# Effect of oral administration of 1,3-1,6 β-glucans in DWV naturally infected newly emerged bees (*Apis mellifera L.*)

**DOI:** 10.1101/2019.12.23.887091

**Authors:** Felicioli Antonio, Forzan Mario, Sagona Simona, D’agostino Paola, Diego Baido, Fronte Baldassare, Mazzei Maurizio

## Abstract

Honeybee pathogens have an important role in honeybee colony mortality and colony losses, most of them are widely spread and necessitate worldwide solutions to contrast honeybee’s decline. Possible accepted solutions to cope with the spread of honeybee’s pathogen are focused on the study of experimental protocols to enhance the insect’s immune defences. Honeybee’s artificial diet capable to stimulate the immune system is a promising field of investigation as ascertained by the introduction of 1,3-1,6 β-glucans as a dietary supplement. In this work, by collecting faecal samples of honeybees exposed to different dietary conditions of 1,3-1,6 β-glucans (0.5% and 2% w/w), it has been possible to investigate the DWV viral load kinetic without harming the insects. Virological data obtained by a one-step TaqMan RT-PCR highlights the ability of 1,3-1,6 β-glucans to reduce the viral load at 24^th^ day of rearing. Furthermore, results indicated that the diet supplemented with 1,3-1,6 β-glucans was associated with a dose-dependent activation of phenoloxidase. Control group showed a higher survival rate than honeybees exposed to different dietary conditions of 1,3-1,6 β-glucans while no differences have been observed concerning syrup daily consumption.

## Introduction

Deformed wing virus (DWV) is one of the most important honeybees’ pathogens and it is responsible for the collapse of the bee colony (de Miranda & Genersch, 2010; Francis et al., 2013). DWV belongs to the *Picornaviridae* family within *Iflavirus* genus (Bailey & Ball, 1991; Lanzi et al., 2006). DWV genome consists of a positive-sense ssRNA of about 7.2 kb (Lanzi et al., 2006). In overt DWV infections, honeybees show malformed or missing wings and shortened abdomens leading to premature death (Kovac & Crailsheim, 1988; de Miranda & Genersch, 2010). Although DWV can be mainly transmitted by the mite *Varroa destructor*, intraspecific and interspecific, vertical and horizontal transmissions have been also demonstrated (Bowen-Walker et al., 1999; Yue & Genersch, 2005; Mazzei et al., 2014; de Miranda & Fries, 2008; Chen et al., 2006b; Yue et al., 2006; Forzan et al., 2017a, 2017b; Mazzei et al., 2018). Several approaches have been proposed for controlling DWV infection (Desai et al., 2012). A previous research indicated that β-glucans, natural molecules constituting the cell wall of many microorganisms (Soltanian et al., 2009; Zecović et al., 2005), have antimicrobial in insects by the activation of *Toll* and *Imd* signalling pathways (Hedengren et al., 1999). The response triggered by β-glucans can be remarkably different depending on the target organism and the dose administered (Guselle et al., 2007; Soltanian et al., 2009). Recent studies demonstrated that high concentration of 1,3 β-glucan are linked to the production of ROS in carp leukocytes and reduction of the immunostimulant effect inducing apoptosis *in vitro* and *in vivo*, (Kepka et al. 2014; Miest & Hoole, 2015). The amount of β-glucans administered is therefore an important factor to be considered to identify the appropriate dosage for the stimulation of the immune response. Moreover, the use of β-glucans, as supplementary food has shown to improve the honeybees’ immune defences (Mazzei et al., 2016). In details, western honeybees (*Apis mellifera L.*) fed with diet based on different 1,3-1,6 β-glucans concentrations have been proved to induce a modification of haemocytes population and an increase of phenoloxidase (PO) activity, a key enzyme involved in pathogen’ encapsulation and nodule formation processes. (Mazzei et al., 2016; González-Santoyo & Córdoba-Aguilar, 2012; Millanta et al., 2019). Furthermore, a 0.5% β-glucans diet was associated with an inhibitory effect on DWV replication (Mazzei et al., 2016).

Since information about the pharmacodynamics action of β-glucans during the experimental period were not previously considered, we planned to measure the DWV viral load from each experimental group by collecting the honeybees’ faeces every three days for 24 days of rearing. Data on diet consumption, survival rate, phenoloxidase and Caspase-3/CPP32 activities were also monitored, recorded and discussed.

## 2. Materials and methods

### 2.1 Rearing conditions

A total of 390 newly emerged worker honeybees were collected in June 2017 from the experimental apiary of the Department of Veterinary Science, University of Pisa, located in San Piero a Grado (PI).

Thirty newly emerged honeybees were collected to constitute T0 group and stocked at −20°C and −80°C for enzymatic and virological assays, respectively. Additional 360 honeybees were sampled and randomly grouped in 12 disposable glass jars (30 individuals each). Honeybees were reared for 24 days in laboratory condition (at controlled temperature 28±2°C in natural dark: light cycle) and fed *ad libitum* with two dosages (0.5 and 2%, w/w) of 1,3-1,6 β-glucans in syrup-based diet and with a constant supply of water.

### 2.2 Feeding conditions

A total of 12 jars with bees were prepared and submitted to three dietary conditions (4 replicates for each condition): syrup only as control diet (G_0_); syrup added with 0.5% (w/w) 1,3-1,6 β-glucans (G_0.5_); and syrup added with 2% (w/w) 1,3-1,6 β-glucans (G_2_). Commercial sugar solution containing 19% glucose, 35% fructose, 12% disaccharides, 6% trisaccharides and 6% polysaccharides was used as control and as 1,3-1,6 β-glucans supplemented diets (Fruttosweet 45 A.D.E.A, Varese, Italy). A fresh mixture of each diet was prepared every three days to avoid any sedimentation of the β-glucans’ content. The feeder contents were checked and refilled every day.

### 2.3 Survival rate measurement and food intake evaluation

To calculate the survival rate, dead honeybees were removed twice a day from each glass jar and counted. Dead individuals were stored at −80°C until further analysis. Each day feeders were weighted before and after refilling to record the food consumption. The weight was normalised to the number of survived honeybees present in the jar.

### 2.4 Sampling

In order to collect honeybees’ faeces, a piece of absorbent paper (0.14 mm thickness, 15.5×10.5 cm) was settled in each glass jar and replaced every three days. The papers collected from each replicate were transferred in disposable plastic bags and immediately processed. Faeces were harvested by making a 1 mm Ø punch on the paper. Twenty punches for each paper were pooled in a 2 ml Eppendorf and stored at −80°C until processed for RNA extraction.

Dead honeybees were collected every day and stored at −80°C until RNA extraction. At day 24^th^ all surviving bees were sacrified, pooled for each experimental group and tested accordingly.

### 2.5 RNA extraction and viral load analysis

Total RNA extraction was performed on faeces, and honeybees using RNeasy mini Kit (Qiagen, Hilden, Germany). RNA from dead honeybees belonging to control and two experimental groups was extracted from pool of three days sampling. After 24 days the survivor honeybees were pooled and RNA extracted to viral load analysis. Briefly, punched papers with faeces or bee samples were homogenized by a stainless-bead (5 mm Ø) in a Tissue Lyser II (Qiagen, Hilden, Germany). Total RNA was eluted in 30 μl RNase-free water and quantified by a Qubit RNA HS kit on the Qubit fluorometer (Thermo fisher scientific, Waltham, MA, USA). Five microliters of each extracted RNA were used as template to determine the viral load by one-step TaqMan RT-PCR assay (Mazzei et al., 2014). Results were expressed as viral copy number per microgram of input RNA for bee sample.

### 2.6 Phenoloxidase activity

Six honeybees from each replicate (24 honeybees per control and experimental groups) were sampled at day 0 and at 24^th^ day of rearing to measure the PO activity. Each bee head was weighed and soaked in 200 µl of 50 mM phosphate buffer pH 7.2 with 1% Triton X-100, left at −20°C for 20 minutes and homogenized by a Teflon pestle. Centrifugation at 4000 rpm, 4°C, for 15 minutes was performed and the supernatant collected. Protein concentration was measured by Qubit Protein Assay Kit on the Qubit fluorometer (Thermo Fisher scientific, Waltham, MA, USA). PO assay was performed by a modified Alaux protocol (Alaux et al., 2010; Mazzei et al., 2016). Absorbance data were obtained at λ=490 nm for 10 minutes. PO values were expressed as UE/mg of tissue.

### 2.7 Caspase-3 activity assay

Caspase-3 activity was measured by the Caspase-3/CPP32 colourimetric Assay Kit (BioVision, USA). The assay is based on the spectrophotometric detection of the chromophore p-nitroaniline (pNA) after cleavage from the labelled N-Acetyl-Asp-Glu-Val-Asp p-nitroanilide substrate (DEVD-pNA). Two pools of three abdomens for each treatment (G_0_, G_0.5_ and G_2_) were tested. Briefly, the abdomens were mixed with 300 μl of PBS buffer pH 7.4, homogenized by a Teflon pestle and, after a macro filtration, centrifuged at 14000 rpm for 4 minutes. Pellets were lysed in 100 μl of lysis buffer and incubated on ice for 10 minutes. After centrifugation at 13000 rpm for 1 minute, supernatants (extracts) were collected and the protein content quantified by Qubit fluorometer (Thermo Fisher scientific, Waltham, MA, USA). The extracts were diluted in lysis buffer in order to obtain the concentration of 1 μg/μl of total proteins. Fifty μl of a 2X Reaction Buffer (containing 10 mM DTT) and 5 µl of the 4 mM DEVD-pNA substrate were added to each extract (50 µl) and incubate at 37°C for 1 hour. Reactions were measured at 405 nm in a microtiter plate reader.

### 2.8 Statistical analysis

Statistical analysis was performed using JMP^©^ software (SAS Institute, 2008). The estimated survival rate was analysed by Wilcoxon rank test using the product-limit (Kaplan-Meier) method for more factors of right-censored data. When factors significantly differed from homogeneous distribution, paired tests were performed. To better highlight the survival rate differences, □^2^ test with Yates correction per day analysis was performed. Food intake, viral load, Caspase-3 assay and PO analysis were performed by the following criteria: data residues obtained by preliminary ANOVA for more factors were tested for normal distribution by Shapiro-Wilk test. When their distribution differed significantly from the normal distribution, these data were analysed by nonparametric Kruskal-Wallis test, while when values were in accord with a normal distribution, the data were analysed by Student t-test.

## Results

### 3.1 Survival rate

Survival rate analysis showed statistical differences among all experimental groups (p<0.001). When survival rates were analysed by a range of 5 days, control group (G_0_) showed higher survival rate than G_2_ from day 6 onwards while resulted higher than G_0.5_ in the 21-24 days interval only (figure 1). At the 24^th^ day of rearing the three experimental groups scored survival values significantly different (p<0.01) with G_2_ (32.5%) and G_0_ (70.3%) groups showing the lowest and the highest survival values, respectively.

**Figure 1.**
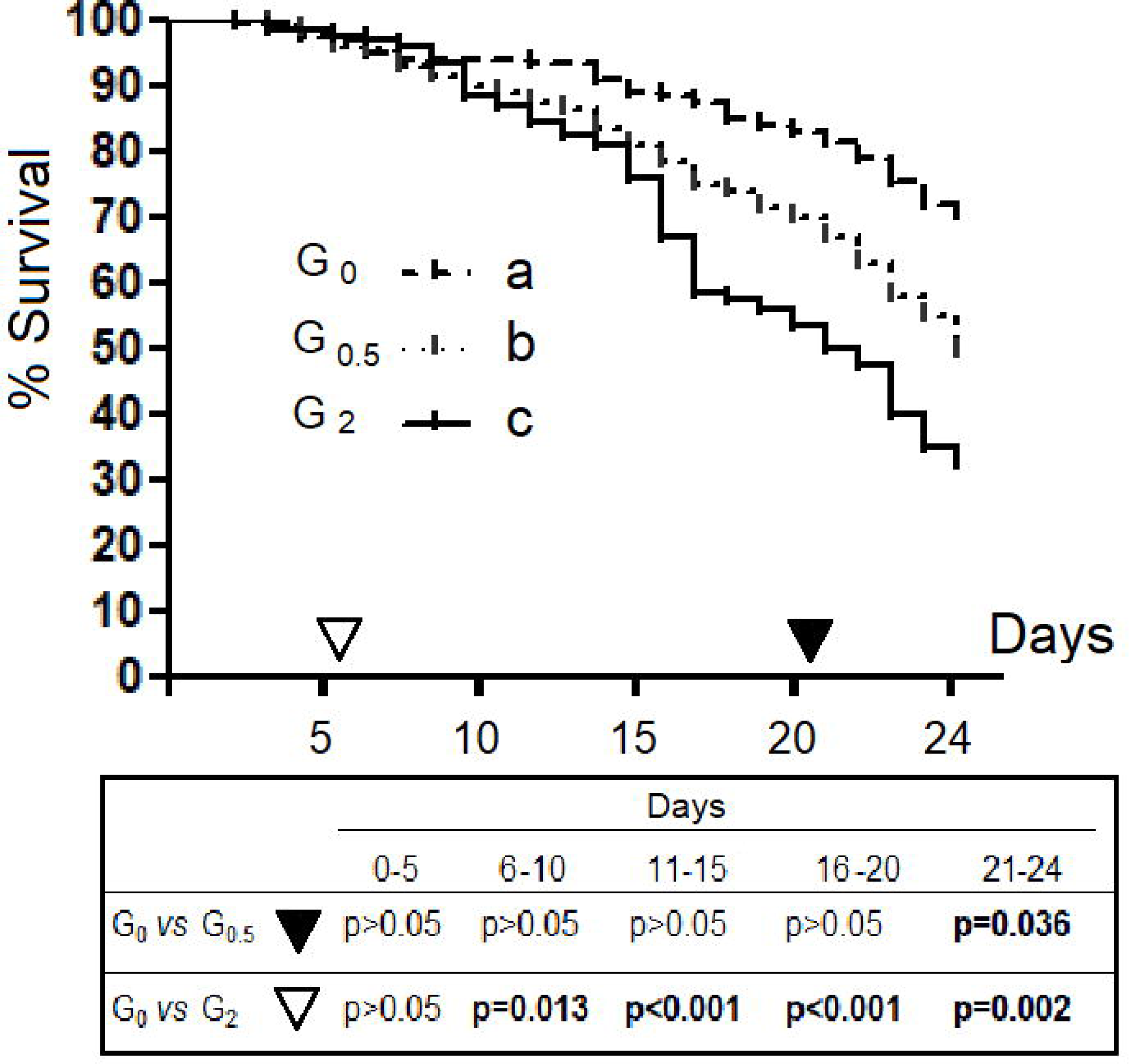
Kaplan-Meier mortality curves: comparison among groups. Each curve indicates the sum of four replicates for group. In G_0_, G_0.5_ and G_2_ are grouped bees feed with 0%, 0.5% and 2% of 1,3-1,6 β-glucans (w/w) in syrup, respectively. Different letters show statistically significant differences (p<0.01).

### 3.2 Food intake

The mean daily syrup consumption for the control and the two experimental groups measured during the whole experimental period resulted in G_0_ 19.9±0.7 mg, G_0.5_ 19.7±0.8 mg and G_2_ 20.9±0.8 mg. No statistical differences were observed among groups.

### 3.3. Viral loads

#### 3.3.1. Viral load of the faecal samples

The kinetics of viral load of the faecal samples of the control and the two experimental groups are reported in figure 2. No significative differences were identified between the two experimental groups and between G_0,5_ and control while only at day 24 G_0_ resulted significantly higher than G_0.5_ (p<0.05).

**Figure 2.**
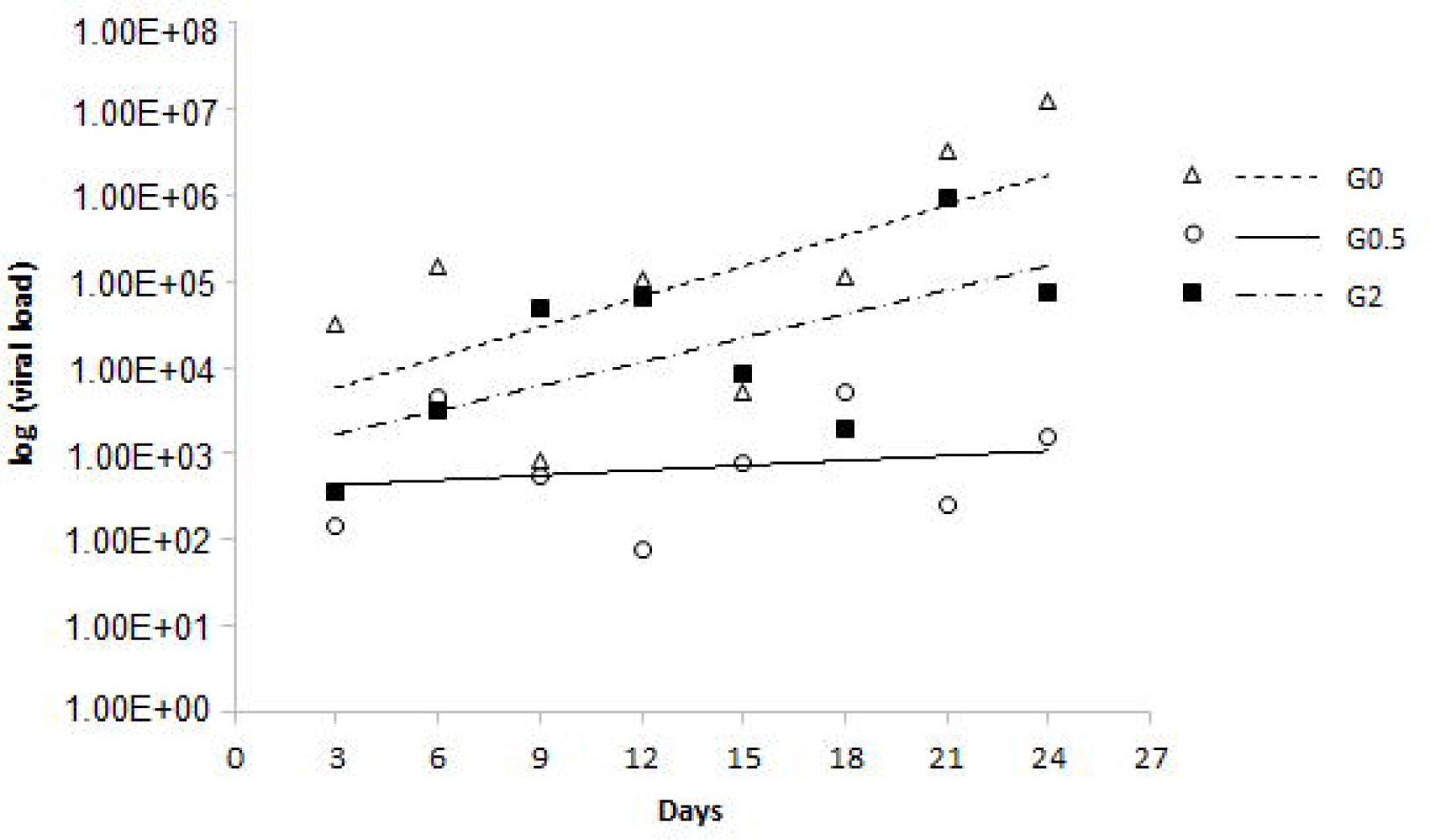
Viral load in the faecal samples from the control and the two experimental groups. Values were expressed as exponential function.

#### 3.3.2. Viral load of the whole dead honeybee samples

No statistical differences were observed for the viral loads of dead honeybees collected during the experiment.

The median for G_0_, G_0.5_ and G_2_ viral load values were 1.05E+05, 1.09E+04 and 8.28E+03, respectively.

#### 3.3.3. Viral load of the whole survived bee samples

The viral load of the survived honeybees analysed at the 25^th^ day, belonging to group G_0_ showed the highest median viral load (4.90E+05/copies DNA microgram), group G_0.5_ an intermediate value (2.22E+05/copies DNA microgram), and the lowest was observed for group G_2_ (5.86E+03), statistically different from the control and the other experimental one, (p<0.01), (fig. 3).

**Figure 3.**
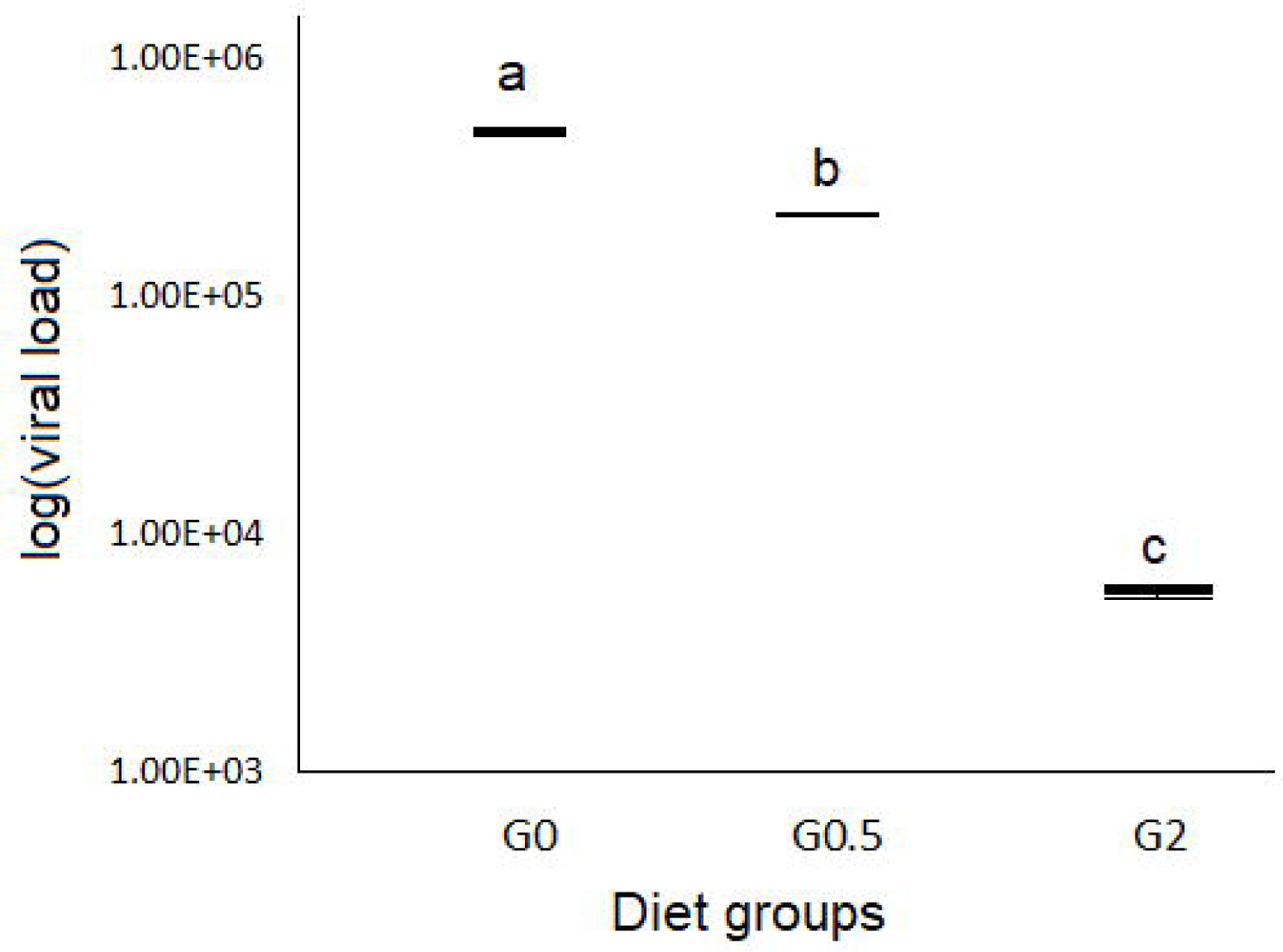
Viral load in the whole insect samples from the three experimental groups. Values were expressed as exponential function. Different letters show statistically significant differences (p<0.05).

### 3.4. Phenoloxidase activity

The PO activity observed at T0 was lower than that observed after 24 days of trial, independently from the diet administered (p<0.01) (figure 4). On day 24^th^, group G_2_ showed the highest PO activity (34.6±1.8 UE/mg of tissue) and G_0_ the lowest (27.2±1.3 UE/mg of tissue) (p<0.01). G_0.5_ showed an intermediate PO value (30.3±1.6 UE/mg of tissue), not significantly different from both the other analysed groups (p>0.05).

**Figure 4.**
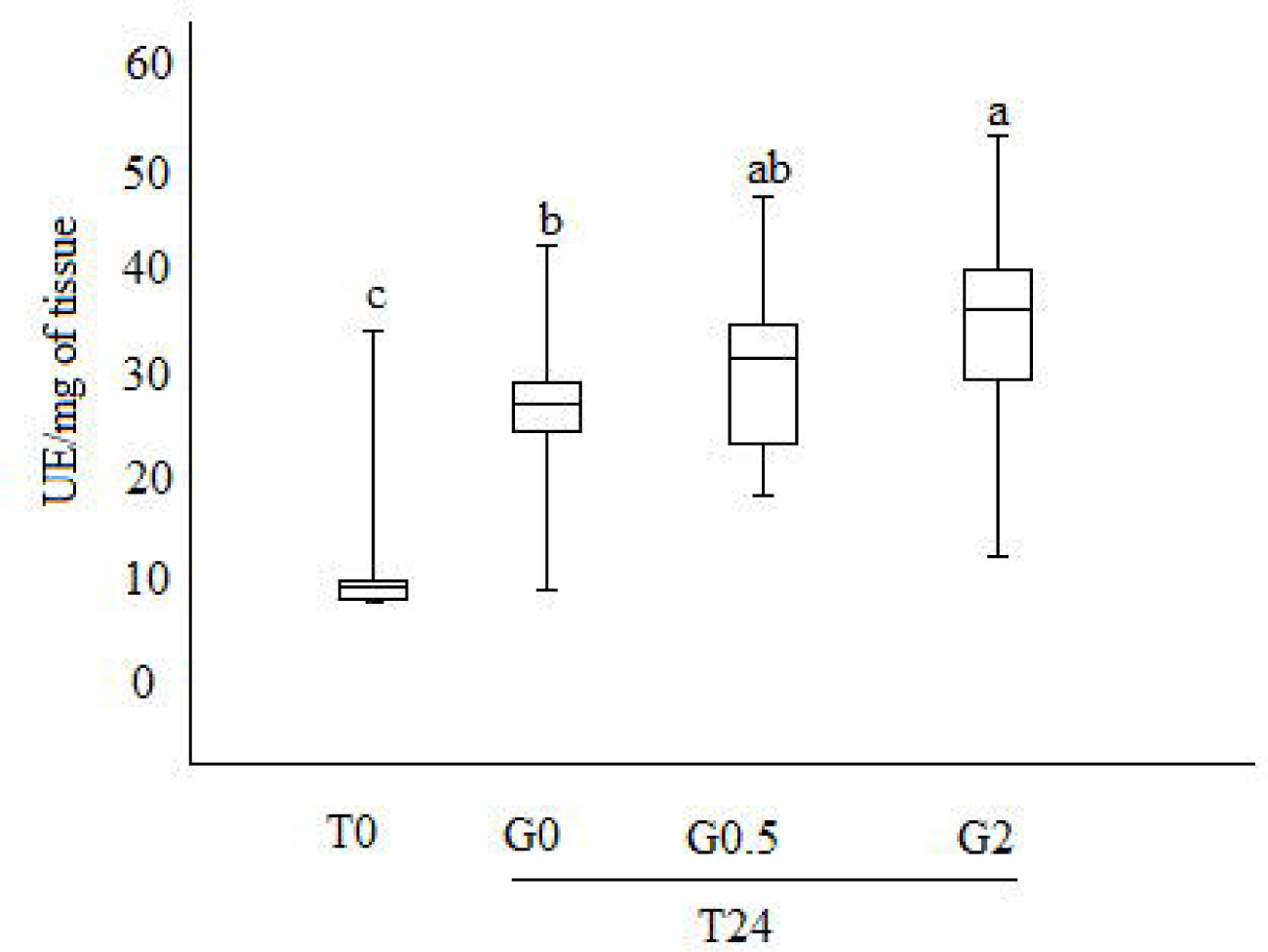
Boxplot representing phenoloxidase activity (expressed in UE/mg of tissue). T0, bees stored at the time of the sample collection; T24, bees after 24 days of rearing; G_0_, G_0.5_ and G_2,_ bees feed with 0%, 0.5% and 2% of 1,3-1,6 β-glucans (w/w) in syrup, respectively. Different letters show statistically significant differences (p<0.01).

### 3.5 Caspase-3 activity assay

The Caspase-3 activity resulted in no significantly different among diet groups. The mean of the absorbance values at 405nm resulted in 0.230±0.036, 0.270±0.023 and 0.168±0.073, for G_0_, G_0.5_ and G_2_, respectively.

## 4. Discussion and conclusion

The use of 1,3-1,6 β-glucans as a diet supplement for DWV infected honeybees has been proven to reduce DWV viral load. Those findings are associated with immune parameters modulation as phenoloxidase activity and haemocytes count (Mazzei et al., 2016). To investigate the effects of a continuous oral administration of 1,3-1,6 β-glucans, honeybees reared in the experimental condition were fed with diets supplemented with different concentration of during a period of 24 days.

The survival rate of honeybees belonging to the control group (G_0_) showed values comparable to those indicated by previous studies performed in honeybees reared in similar conditions (Di Pasquale et al., 2013; Frias et al., 2016). Results obtained in this investigation indicate that the presence of 1,3-1,6 β-glucans in the syrup diet had a negative dose-dependent effect on the survival rate. The diet containing 2% and 0.5% β-glucans supplement caused a lower survival rate in comparison to control diet. The low survival rate recorded for G_2_ and G_0.5_ group is in consequence of increased mortality recorded in the 6-10 and 21-24 days intervals, respectively. This result suggests a possible dose-dependent toxic effect due to the β-glucans administration, that could induce a prolonged stimulation of immune defence system, resulting in a negative effect of metabolic intermediates accumulation as previously reported in invertebrate and in insect by Söderhäll & Cerenius (1998) and González□Santoyo & Córdoba□Aguilar(2012), respectively.

No statical differences were observed on the food intake among control bees and the two experimental groups, indicating that 1,3-1,6 β-glucans does not determine lower or higher attractiveness than only syrup diet. Caspase-3 activity was not different among groups. Since caspase is an indicator of apoptosis this result indicated that β-glucans administrated as a supplementary diet on honeybees does not induce apoptosis.

A significant difference in viral loads in faecal samples was detected at the 24^th^ day only between G_0_ and G_0.5_. At the best of our knowledge, since faecal excretion of the virus from infected honeybees has been previously reported (Chen et al., 2006b), this is the first report in which the kinetic of DWV infection in honeybees faeces has been monitored by an RT-qPCR. The results obtained in this investigation indicate the RT-qPCR on faeces samples as a useful diagnostic tool to quantify DWV in living honeybees.

No statistical differences were observed for the viral loads of dead honeybees collected during the course of the experiment. Noteworthy, viral load of survived honeybees belonging to G_0.5_ and G_2_ was lower than the control group. This result strongly suggests a relationship between 1,3-1,6 β-glucans and DWV. The highest values of PO at day 24th scored by honeybees fed with 2% β-glucans could be due to the well-known ability of 1,3-1,6 β-glucans to activate the proteases involved in activating phenoloxidase, as reported by Vetvicka & Sima (2004) in invertebrates, which, in turn, is involved in melanisation of pathogens.

When compared viral load values with PO activity on the surviving bees, G_0_ group scored the highest viral load and the lowest PO activity while G_2_ showed the lowest viral load and the highest PO activity. G_0.5_ showed intermediate values for both parameters. Such results could be interpreted as a 1,3-1,6 β-glucans dose dependent activation of PO that results in a viral replication restraint.

This research confirmed 1,3-1,6 β-glucans as molecules able to modulate honeybees defence pathways. To better characterize and define the 1,3-1,6 β-glucans effects on honeybees, further evidences should be necessary in order to define their posology and method of administration.

## Supporting information

Cover letter

