## Supplementary material for "Effect of oral administration of 1,3-1,6 β-glucans in DWV naturally infected newly emerged bees (*Apis mellifera L.*)": Cover letter

Maurizio Mazzei

Department of Veterinary Science

University of Pisa

Viale delle Piagge, 2

56124 PISA – ITALY

December 23, 2019

Dear Editor,

I am pleased to submit an original research article “Effect of oral administration of 1,3-1,6 β-glucans in Deformed Wing Virus naturally infected newly emerged bees (*Apis mellifera L.*)” for consideration for publication in Journal of Apicultural Research. In this manuscript, the effect of 1,3-1,6 β-glucans on DWV viral load kinetic, collecting faecal samples of honeybees exposed to different dietary conditions has been investigated without harming the insects.

We believe that this manuscript is appropriate for publication by in Journal of Apicultural Research. Our manuscript confirms 1,3-1,6 β-glucans as molecules able to modulate honeybees defence pathways, moreover indicate the RT-qPCR on faeces samples as a useful diagnostic tool to quantify DWV in living honeybees.

This manuscript has not been published and is not under consideration for publication elsewhere. We have no conflicts of interest to disclose.

Thank you for your consideration!

Sincerely,

Maurizio Mazzei PhD, DVM

Associate Professor, Department of Veterinary Science

University of Pisa

Authors details:

Felicioli Antonio	https://orcid.org/0000-0002-0808-3037,

Forzan Mario	https://orcid.org/0000-0003-2474-8219

Sagona Simona	https://orcid.org/0000-0003-1592-7522,

D’agostino Paola

Diego Baido

Fronte Baldassare	https://orcid.org/0000-0002-8480-318X,

Mazzei Maurizio	https://orcid.org/0000-0003-3797-8320
